# Plants sum and subtract stimuli over different timescales

**DOI:** 10.1101/2023.01.06.522981

**Authors:** Mathieu Rivière, Yasmine Meroz

**Affiliations:** Faculty of Life Sciences, School of Plant Science and Food Security, Tel Aviv University, Tel Aviv, Israel

## Abstract

Mounting evidence suggests that plants engage complex computational processes to quantify and integrate sensory information over time, enabling remarkable adaptive growth strategies. However, quantitative understanding of these computational processes is limited. We report experiments probing the dependence of gravitropic responses of wheat coleoptiles on previous stimuli. First, building on a mathematical model that identifies this dependence as a form of memory, or a filter, we use experimental observations to reveal the mathematical principles of how coleoptiles integrate multiple stimuli over time. Next, we perform two-stimulus experiments, informed by model predictions, to reveal fundamental computational processes. We quantitatively show that coleoptiles respond not only to sums but also to differences between stimuli over different timescales, constituting first evidence that plants can compare stimuli – crucial for search and regulation processes. These timescales also coincide with oscillations observed in gravitropic responses of wheat coleoptiles, suggesting shoots may combine memory and movement in order to enhance posture control and sensing capabilities.

## Introduction

Plants, decentralized systems, negotiate fluctuating environments by adapting their morphology advantageously through growth-driven processes called tropisms, whereby organs such as shoots or roots redirect their growth according to directional stimuli, such as gravity or light. The last decade has seen an increasing body of evidence that plants quantify and integrate sensory information about their environment at the tissue level (1–16), detecting and climbing spatial signal gradients despite environmental and internally-driven noise (17). While it is generally accepted that tropic responses are a product of complex computational processes (1, 14, 18–22), attempts to characterize these processes mathematically— that is, to obtain an understanding of the rules according to which plants quantify and process the sensory information they acquire over time—are limited. There is a need for comprehensive experimental and mathematical frameworks that can be used, respectively, to derive quantitative observations regarding plants’ growth in response to various stimuli, and to decode the underlying principles.

Here, we report experiments that probe the dependence of gravitropic responses of wheat coleoptiles on the presence of previous stimuli, building on previous findings that plant shoots respond to the integrated history of stimuli rather than responding instantaneously (13, 23–29). Our work comprises two phases. First, we subject coleoptiles to a single gravita-tional stimulus, of varying duration, and record their gravitropic response over time. From these observations we extract a functional representation of the collection of computational processes underlying plants’ gravitropic responses. This phase relies on a mathematical model that was previously developed to describe temporal integration in plant tropisms, grounded in *response theory* or *control theory* (see below) (12, 13). In the second phase of this study, on the basis of the characteristic shape of the extracted function, we make predictions (supported by model simulations) regarding how plants integrate multiple stimuli over different timescales. We then conduct two-stimulus experiments which corroborate our predictions. These results provide the first quantitative evidence that coleoptiles effectively respond not only to the sums of stimuli, as previously reported (13, 23–29), but also to the differences between stimuli over different timescales. The latter suggests that plants have the ability to compare signals—a critical capability in search processes.

## Results

subsection*Model description Fig. 1a shows an example of the evolution of the gravitropic response of a wheat coleoptile tilted horizontally on a platform, redirecting its growth in the direction of gravity. For a given snapshot at time *t* we represent the coleoptile’s geometrical shape relative to the frame of reference of the platform, described in Fig. 1a. The shape is defined by (i) the local angle from the vertical *θ*(*s, t*) at point *s* along the organ, where *s* = 0 at the base and *s* = *L* at the tip, and (ii) the local curvature 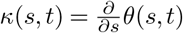, which is the rate of change of the angle along the organ. We denote the angle at the base *θ*_base_, the angle at the tip *θ*_tip_, and the angle of the stimulus *θ*_P_, here the direction of the platform relative to gravity.

**Fig. 1.**
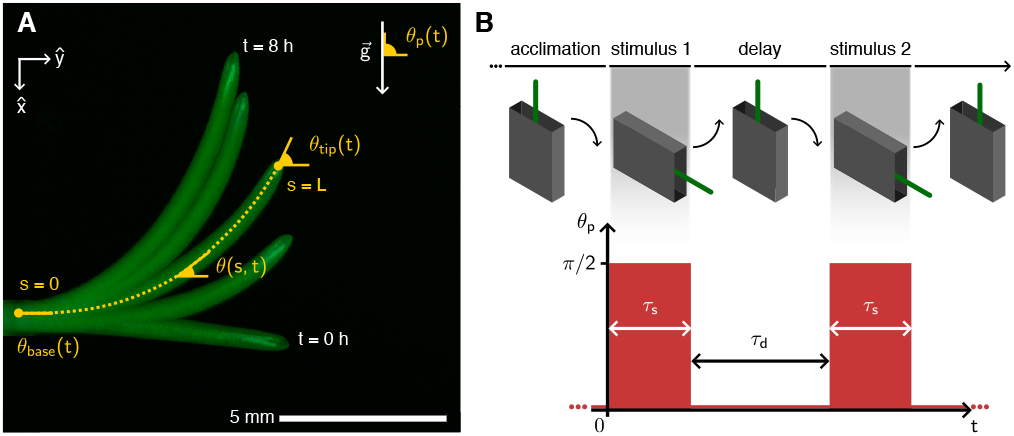
Transient gravistimulation of wheat coleoptiles. **(A)** Typical gravitropic response of a wheat coleoptile to an inclination of *θ*_P_ = *π/*2 with respect to the direction of gravity 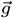 (snapshots taken every 2 hrs). After 6 hrs the tip is aligned with the direction of gravity. Coleoptiles are described in the frame of the rotating platform (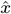, *ŷ*). The arclength *s* spans the whole midline, with *s* = 0 at the base and *s* = *L* at the tip, where *L* is the organ length. The local angle relative to *ŷ* is given by *θ*(*s, t*). In addition, we denote the angle at the base *θ*(*s* = 0, *t*) = *θ*_base_, and the angle at the tip *θ*(*s* = *L, t*) = *θ*_tip_. **(B)** Sketch of the experimental protocol. Coleoptiles are attached to the rotating platform and are either tilted once (*θ*_P_ = *π/*2) for a duration *τ*_s_ and then tilted back to vertical position, or receive two successive stimuli of equal duration *τ*_s_ delayed by *τ*_d_.

The mathematical model at the basis of our approach (12, 13) incorporates concepts from *response theory* or *control theory*, commonly used in the description of input-output systems, signal transducers (30), and dynamics responses of sensory-motor systems (31). In its simplest linear form, the response *y*(*t*) of a system is described as a weighted integral of the history of stimuli *x*(*t*) over time, where the weight is given by the response function, or memory kernel, *μ*(*t*), following

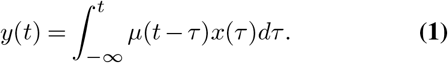

In simple cases where the dynamical equation describing an input-output system is known, the solution to the equation can be rewritten in the form described in Eq. 1, and the memory kernel is directly related to the original equation (30). Response theory is however particularly useful for describing systems in which the dynamical equations are not known, such as complex biological systems (32–35). In principle, the form of the memory kernel in such formulations is a mathematical representation of underlying equations, a transfer function, agnostic of the building blocks of the system. In terms of signal processing or control theory, the memory kernel corresponds to a filter (31).

Approaching the case of plant gravitropism through the lens of response theory, we identify the dependence of gravitropic responses on past stimuli as a form of memory, captured by the memory kernel. Examples of memory formation at the tissue level are widespread (18, 21, 22, 36, 37), even in non-biological matter (38, 39). Here we assume that the input signal depends on the relative angle between the organ and the direction of gravity, following the well-known sine law sin(*θ*(*s, t*) −*θ*_P_) (40, 41). The signal is zero if the organ is aligned with gravity, and maximal if it is perpendicular to it. Following Eq. 1 we convolute the signal with the memory kernel *μ*(*t*), yielding the transduced signal as an output, linear with respect to the signal. This linearity suggests that the response of the system should abide by the superposition principle of two stimuli, which we verify in Fig. 3. This transduced signal, in turn, dictates the tropic dynamics following (12):

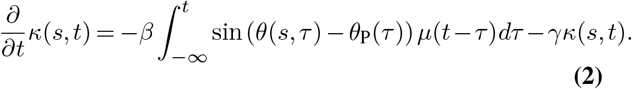

The tropic dynamics are represented by the change of curvature in time *∂κ*(*s, t*)*/∂t*, and is driven by the transduced signal due to a varying inclination angle *θ*_P_(*t*), and a relaxation term *γκ*(*s, t*) associated with *proprioception* – the tendency of a plant to grow straight in the absence of any external stimuli (42). The coefficients *β* and *γ* are, respectively, the gravitropic and proprioceptive sensitivities or gains, which vary across species (42). Growth is considered as the implicit driver of the tropic response, and is not explicitly taken into account. Eq. 2 therefore holds within the growth zone of length *L*_gz_ from the tip. In the mature zone, where no growth-driven bending can occur, the equation reduces to 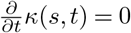.

Here, the memory kernel represents the dynamics of a hierarchy of stochastic processes underpinning gravitropic responses (12, 13, 19), the details of which are currently not well understood (43), such as statolith sedimentation in gravity-sensing cells, and the ensuing asymmetrical redistribution of PIN proteins and the growth hormone auxin (44). Previous studies adopting a *Response Theory* approach to model tropic responses assumed an arbitrary form of the memory kernel, and showed that the model qualitatively reproduced observations of temporal integration of multiple stimuli over limited timescales (12, 13), however could not make quantitative predictions, or explain negative responses observed for transient stimuli at longer times (13). To obtain a quantitative picture of plants’ computational capabilities, in terms of quantifying and processing sensory information, it is necessary to extract the mathematical form of the memory kernel.

### Estimation of the memory kernel

Accordingly, in the first phase of this study, we sought to extract the memory kernel by probing the dependence of gravitropic responses on the history of stimuli. To this end, we exposed wheat coleoptiles to transient gravistimulation protocols. Coleoptiles were placed vertically on a platform, which inclined horizontally for a stimulus duration *τ*_s_, then rotated back to the vertical (shown schematically in Fig. 1b). We recorded the gravitropic responses of coleoptiles to different values of stimulus duration, ranging from *τ*_*s*_ = 6 min, the shortest to elicit a clear response, to a permanent stimulus *τ*_*s*_ → ∞ (in practice, *τ*_*s*_ = 10 hours). The average gravitropic response, represented by the average evolution of the tip angle *θ*_tip_(*t*), are shown in Fig. 2a for the limiting cases of *τ*_*s*_ = 6 and *τ*_*s*_ *→ ∞*. Fig. 2b shows the normalized response 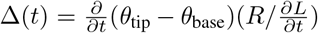 (45) for the same two examples. The normalized response accounts for variations in growth rate 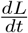 and organ radius R, allowing us to better compare experimental trajectories to simulations later on. For each response trajectory, of a single coleoptile, we numerically solved Eq. 2, and extracted the mathematical form of the memory kernel (elaborated in the Methods), substituting a low-dimensional representation of the organ shape as measured by the values of *θ*(*s, t*) and *κ*(*s, t*), and the stimulus profile with *θ*_P_(*t*) = *π/*2 during the stimulus duration *τ*_*s*_, and *θ*_P_(*t*) = 0 otherwise. For each stimulus duration *τ*_*s*_ we averaged over the extracted memory kernels, and the average kernels 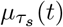 are shown in Fig. 2c.

**Fig. 2.**
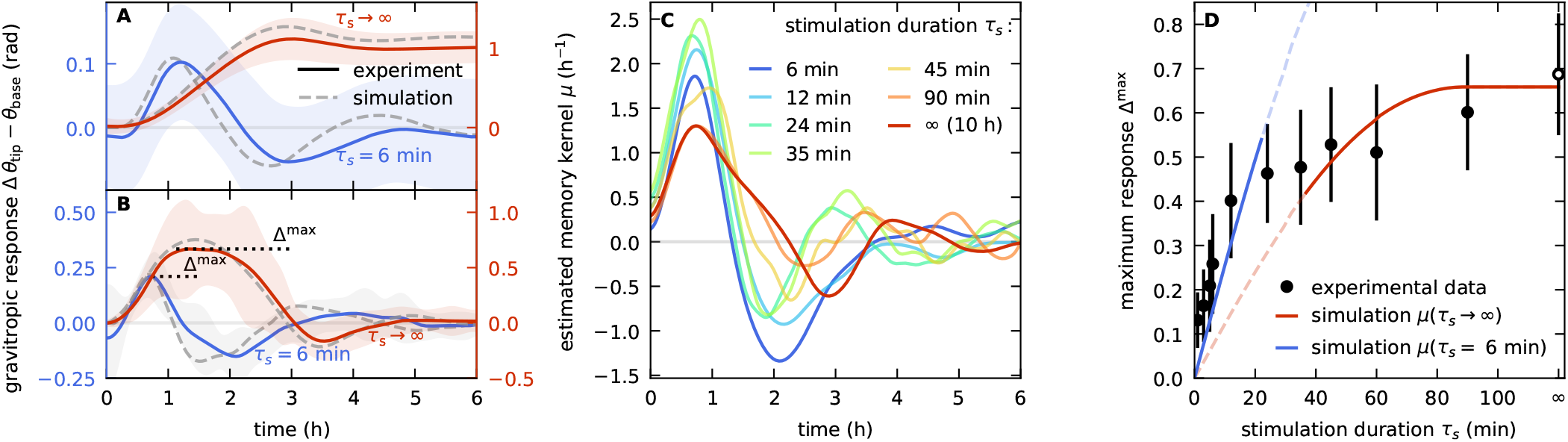
Estimated memory kernels are consistent with experiments and capture the dynamics of gravitropism. **(A)**The average tip angle trajectory *θ*_*tip*_(*t*) − *θ*_*base*_(*t*) of coleoptiles inclined for a short stimulus time *τ*_s_ = 6 *min* (solid blue) and permanent stimulus *τ*_s_ → ∞ (solid red). Shaded areas represent standard deviation. Simulated tip dynamics (dashed lines), based on average memory kernels estimated from the respective data, *μ*_6_ and *μ*_*∞*_, agree with experimental data. **(B)** Similar comparison for the normalized gravitropic response Δ(*t*) = *R∂*_*t*_(*θ*_*tip*_ *−θ*_*base*_)*/∂*_*t*_*L*. Δ^max^ is the maximal response after the stimulus. **(C)** Averaged memory kernels extracted from experiments with various values of *τ*_s_. **(D)**. Maximum gravitropic response Δ^max^ for increasing stimulus duration *τ*_s_. Values increase for short stimulus times, and saturate at longer times. Each experimental value (black dots) was obtained from an average with *N* ≥ 30 repetitions, errors represent standard deviation. The empty dot represents a permanent stimulus. Simulations based on the limiting memory kernels extracted from responses to the shortest and longest stimulus times, *τ*_s_ = 6 *min* (blue) and *τ*_s_ → ∞ (red), capture both regimes.

In order to verify our model accurately captures the average response over time, we simulated the gravitropic response for each *τ*_*s*_, based on the corresponding average memory kernel 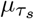. We compared the simulations to the average tip angle responses and normalized responses. These were found to be in excellent agreement, as shown in the examples in Fig. 2a-b. Fig. S7 shows kymographs, or heatmaps, displaying the evolution of the local curvature *κ*(*s, t*) along the whole shoot for representative experiments for *τ*_s_ = 6 min as well as *τ*_s_ *→ ∞*.

While the heatmap for the permanent stimulus recovers previous measurements (42), the heatmap for *τ*_s_ = 6 min exhibits complex local dynamics. In the SM we show that the response is robust to small perturbations in the extracted graviceptive and proprioceptive gains *β* and *γ* (Fig. S1), as well as perturbations in *τ*_*s*_ (Fig. S2).

### Memory kernel characteristics and model validation

We observe that all memory kernels extracted from responses to different *τ*_s_ (Fig. 2c), exhibit similar characteristics; a positive peak followed by a negative peak, which then goes to zero. The timescales and magnitudes of the two peaks differ slightly across kernels (detailed in the SM), suggesting that the memory kernel probes different underlying biological processes. This is in line with arguments tying the dependence of gravitropic responses on stimulus duration (13) to the different timescales of underlying processes, from statolith sedimentation, dynamics of PIN proteins and auxin transport, and the ensuing growth response (43). Following Chauvet et al. (13), we plotted the maximal values of the average normalized responses Δmax = max {Δ(*t*)} for different *τ*_*s*_ (Fig. 2d). We recovered two regimes, namely, increasing responses for short *τ*_*s*_, and saturation at long *τ*_*s*_ (13). We examined the dependence of the gravitropic responses on the form of the memory kernels, finding that simulations of responses to the range of *τ*_s_, using only the limiting memory kernels *μ*_6_ and *μ*∞ extracted from the responses to *τ*_s_ = 6 min and *τ*_s_ = ∞, recovered the behavior in both regimes (Fig. 2d). This result further corroborates our model and the extracted memory kernels.

### Validation of kernel linearity

We assess the validity of the assumption of a linear response in time, expressed by the convolution of the signal with a memory kernel. We examined coleoptiles’ gravitropic responses to two consecutive stimuli of *τ*_*s*_ = 6 min each, separated by a delay time *τ*_d_ (Fig. 1B). For clarity, in what follows we will denote the responses to single stimuli or double stimuli as Δ_1_(*t*) and Δ_2_(*t*) respectively. In a linear system we expect the measured response to two stimuli, Δ_2_(*t*), to be equal to the sum of the response to a single stimulus Δ_1_(*t*), and the response shifted by *τ*_d_, Δ_1+1_(*t*) = Δ_1_(*t*) +Δ_1_(*t −τ*_d_). Fig. 3A shows a specific example, comparing Δ_2_(*t*) for stimuli separated by *τ*_d_ = 90 min (black solid line), with the predicted response Δ_1+1_(*t*), the sum of individual responses to a single stimulus shifted by *τ*_d_ = 90 min (black dashed line). The two are in very good agreement. In order to quantify this, we define the maximal values of the measured response Δ_2_(*t*) and predicted response Δ_1+1_(*t*) after the second stimulus,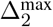 and 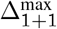 respectively, as well as their time of occurrence, 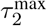 and 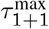. Fig. 3B plots the measured maximal values 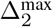. the predicted values 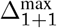 for a range of waiting times *τ*_d_. Fig. 3C plots 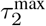 vs. the predicted times,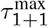. The predicted values are in very good agreement with observations, confirming the model assumption that the system is linear. A similar analysis was carried our for the memory model described in Eq. 2, running simulations for a range of stimulus times *τ*_s_ and waiting times *τ*_d_. Results are shown in Fig. 3D, finding that the model is indeed linear for *τ*_s_ = 6 min, as is the case of the experimental observations. However linearity weakens as *τ*_s_ increases.

**Fig. 3.**
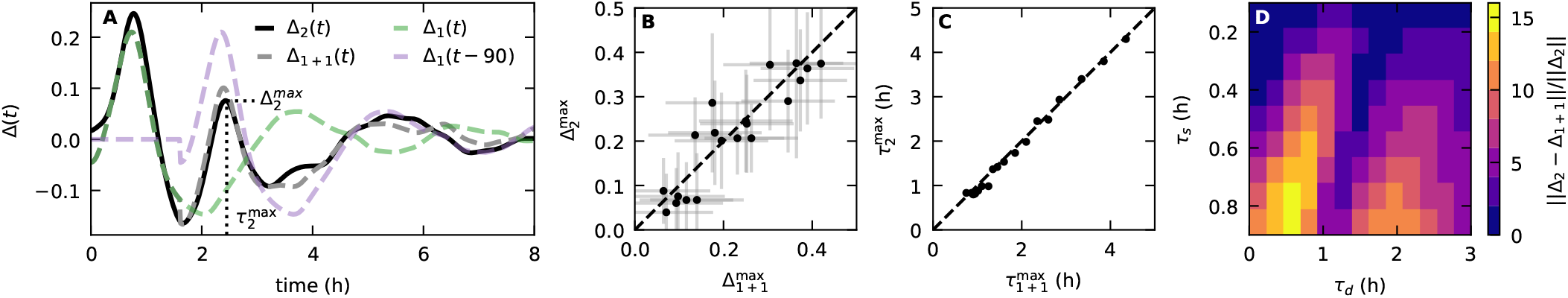
Validation of temporal linearity. **(A)** Example of the response Δ_2_ (*t*) measured for two stimuli of duration *τ*_s_ = 6 min separated by *τ*_d_ = 90 min (solid black), in excellent agreement with the predicted response for a linear system Δ_1+1_ = Δ_1_ (*t*) + Δ_1_ (*t −*90) (dashed black line), where Δ_1_ (*t*) is the response measured for a single stimulus (green), and Δ_1_ (*t −*90) os the response shifted by the delay time (purple). The two are in excellent agreement. We represent the responses by the maximal value following the second stimulus,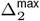 and 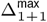, and their occurrence,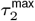 and 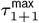 respectively, marked by a dashed line. **(B)** Comparison of measured and predicted maximal responses,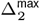 and 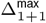, for two-stimulus experiments with *τ*_d_ values ranging 1 − 240 min, values detailed in the Methods. Dashed line represents identity. **(C)** Occurrences of the measured and predicted responses,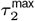 and 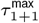, for all two-stimulus experiments. Dashed line represents identity. **(D)** Linearity assessment of the mathematical model (Eq. 2). The x-axis represents different waiting time *τ*_d_, the y-axis represents different stimulus durations *τ*_s_. For each combination we ran a simulation and calculated Δ_2_ and Δ_1+1_, and the color-code corresponds to the distance between the functions in % relative to Δ_2_. The top row represents the experimental system with *τ*_s_ = 6 min, exhibiting small distances (smaller than 3%) suggesting the system is linear. At larger *τ*_s_ distances increase.

### Memory kernel reveals computational processes

In the second phase of our study, we sought to gain an understanding of the specific computational processes represented by the characteristic form of the extracted memory kernel. To this end, we examined the responses to single and double stimuli, Δ_1_(*t*) and Δ_2_(*t*), described before and in the Methods (Fig. 3). To provide intuition as to the patterns we might expect to see for different values of *τ*_d_, and the computational processes they reveal, Figs. 4A–C show schematics of stimuli with different timescales overlaid on a characteristic memory kernel. For short *τ*_d_, both stimuli fit within the first positive peak, and are both weighted positively (Fig. 4A). The coleoptile is therefore expected to respond to the weighted *sum* of the two stimuli; i.e., the response to the second stimulus is expected to be stronger than the response to a single stimulus. Fig. 4D shows the average experimental response Δ_2_(*t*) for *τ*_d_ = 10 min, well within the timescale of the first peak (compare to *μ*_6_ in Fig. 2C). For reference, the two stimuli are marked with grey bars, as well as the maximal response 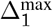 for a single stimulus *τ*_s_ = 6 min represented by a vertical line (taken from Fig. 2B). The maximal value 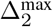 after the second stimulus is significantly higher than the response to a single stimulus 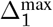 (illustrated by an arrow), verifying the prediction of our model. For intermediate *τ*_d_, the second stimulus occurs within the positive peak and the first stimulus within the negative peak, and is therefore multiplied by a negative value (Fig. 4B). The coleoptile is therefore expected to respond to the weighted *difference* of the two stimuli; i.e., the response to the second stimulus is expected to be weaker than the response to a single stimulus. This prediction is experimentally verified for *τ*_d_ = 90 min (Fig. 4E). Finally, for *τ*_d_ longer than the timescale of *μ*(*t*), while the second stimulus occurs within the positive peak, the first stimulus is multiplied by a null value of *μ*(*t*) (Fig. 4C), and is effectively *forgotten*. The response to the second stimulus is expected to be similar to that of a single stimulus, corroborated in Fig. 4F for *τ*_d_ = 10 hr.

**Fig. 4.**
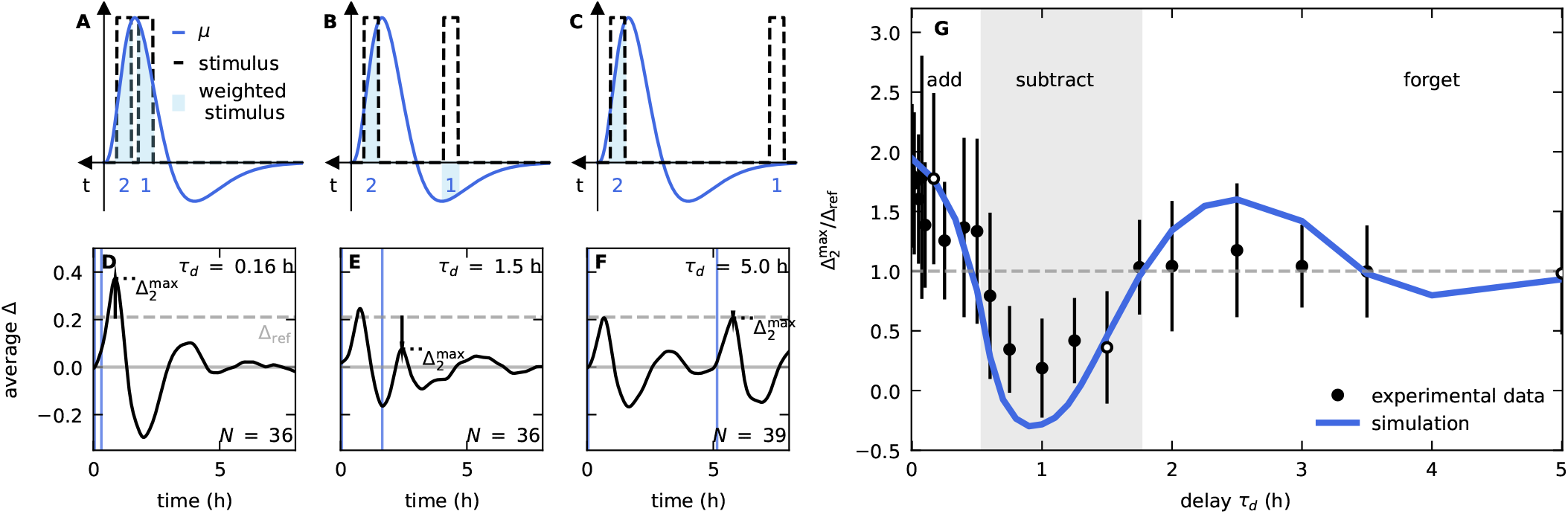
Model prediction and experimental observations of summation and subtraction regimes in two-stimulus experiments. **(A–C)** Graphical illustration of *μ* (blue line) compared to two stimuli (dashed bars) separated by different delay times *τ*_d_. Shadowed areas correspond to stimuli weighted by *μ*. **(A)** For short *τ*_d_, both stimuli occur within first peak, and are weighted positively. Coleoptiles are expected to respond to the weighted sum of stimuli, with a stronger response than for a single stimulus: verified in **(D)** the experimental response Δ_2_ (*t*) for *τ*_d_ = 10 min (compare to *μ*_6_ in Fig. 1C). For reference, the dashed line represents 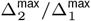 for single stimulus. **(B)** For intermediate *τ*_d_, second stimulus occurs within the positive peak, weighted positively, and the first within the negative peak, weighted negatively. Coleoptiles expected to respond to weighted *difference* between stimuli, with a weaker response compared to a single stimulus: verified in **(E)**, the response for *τ*_d_ = 90 min, roughly the time between the two peaks in *μ*_6_. **(C)** For *τ*_d_ larger than the timescale of *μ*, the second stimulus occurs within the positive peak, weighted positively, while the second is multiplied by zero. The response is expected to be similar to that for a single stimulus: verified in **(F)**, for *τ*_d_ = 5 hrs. **(G)** Maximal response of two-stimulus experiments relative to single stimulus 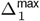, for a range of *τ*_d_. Circles represent experimental data, and the blue line represents the simulated response based on the extracted kernel *μ*_6_. Shadowed area delineates the three regimes predicted in **(A—C)**. The three white circles represent experiments **(D—F**).

We extended these two-stimulus experiments, in addition to simulations, for a range of values of *τ*_d_ between 0 and 5 hours, spanning the timescale of *μ*_6_. In Fig. 4G we report the values of the maximal response 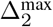 following the second stimulus, relative to 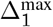 as a reference, as a function of the delay time *τ*_d_, both for experiments and simulations. Our two-stimulus experiments, supported by simulations, reveal the three regimes we predicted earlier based on the form of the extracted memory kernel, illustrated in Fig. 4A–C: (i) for short *τ*_d_ coleoptiles respond to the weighted sum of stimuli, in line with previous observations of temporal integra-tion (12, 13, 25–29), and the response to the recent stimulus is stronger than for a single stimulus; (ii) for *τ*_d_ of the order of the time between peaks, coleoptiles respond to the weighted difference of stimuli, and the response is weaker; and (iii) for longer *τ*_d_ the coleoptiles respond to the recent stimulus alone. In the SM we show that these results are not an artifact of the relative orientation of the organs at the time of the second stimulus (Fig. S3). Furthermore, while it is possible that proprioception may have a role in the observed memory, we note that it is a second-order response to the stimulus, mediated by the curvature. We also show that a neglected kernel in the proprioceptive term would not significantly affect the extracted gravitropic kernel for similar timescales (Fig. S4).

## Conclusions

Together, these findings provide the first quantitative evidence that plants respond to the sum and *difference* of stimuli over different timescales, framed within a general mathematical framework. These computational processes can be identified as fundamental elements of natural search algorithms, common across diverse biological organisms (46). In the context of signal processing or control theory, the memory kernel may be interpreted as a band-pass filter. The summation of stimuli is effectively a moving average, improving the signal-to-noise ratio of an environmental signal. Subtraction of stimuli, which has never been reported before in plants, is required in order to compare signals over time, a strategy commonly used by a variety of living organisms in order to detect and climb signal gradients (47). For example, Segall et al. (33, 48) extracted a memory kernel describing the chemotactic response of bacteria to a chemical stimulus, and found that it, too, was characterized by a positive peak followed by a negative one. On this basis they found that bacteria compare chemical concentrations sampled over time, enabling them to identify chemical gradients.

Observed oscillations in the gravitropic reorientation of wheat coleoptiles have been conjectured to speed up the regulation of posture control (14, 49, 50). The half period of observed oscillations was found to be 1.1 hr (49, 50), consistent with the delay time *τ*_*d*_ of maximal subtraction, and anything occurring further than a full oscillation period (2.2 hr) is *forgotten* (Fig. 4G). Within this context, our findings suggest that shoots may enhance regulation of posture control by comparing the relative inclination of the organ sensed at either side of an oscillation or circumnutation period, or equivalently detect a light gradient by comparing measured light intensities. Furthermore, as the ability of plants to integrate stimuli has been observed for a range of species and tropisms, it suggests that these computational abilities might be general.

Put together, our study provides the backdrop for understanding how plants may combine memory and movement in order to enhance movement control and sensing capabilities in the face of weak signals and fluctuating environments.

## Materials and methods

### Wheat growth conditions

Wheat coleoptiles were grown as described in (13, 45). Wild-type wheat seeds (*Triticum aestivum* cv. Ruta) were initially glued with floral glue to plastic boxes with draining holes at the bottom (germ pointing down). The boxes were filled with cotton and placed in a light-proof container, immersed in water. Coleoptiles were kept in a growth chamber at a temperature of 24 °C for 3 to 4 days. Coleoptiles were manipulated in darkness or dim green light to minimize exposure to actinic light.

### Experimental setup

Our experimental apparatus consisted of an upright rotating platform holding 3D-printed casings each hosting a coleoptile box. A DC motor inclined the platform, and the inclination angle was measured and controlled with two optical fork sensors and a custom 3D-printed encoder wheel. During experiments images were taken every 1, 5 or 10 min, depending on the experiment, using a DSLR camera with a wide-angle lens, controlled by the Arduino. Pictures were taken with a dim green flash of 333 ms, placed perpendicular to the plane of motion. The setup was controlled by an Arduino microcontroller board running an in-house developed program. Each experiment started by placing upright coleoptile boxes on the rotating platform, and waiting for an acclimation phase of 10 h in the dark. Coleoptiles were then gravistimulated by rotating the platform horizontally, *θ*_*p*_ = *π/*2, during a stimulation time *τ*_s_, and then rotated back to the vertical position. We employed two types of protocols (schematically illustrated in Fig. 1B): (i) single stimulus, for a range of *τ*_s_ values (in minutes: 1, 3, 5, 6, 12, 24, 35, 45, 60, 90 and 600), (ii) double stimulus; apply two successive stimuli of *τ*_s_ = 6 min each, separated by a delay time *τ*_d_ (in minutes: 1, 3, 4.5, 6, 10, 15, 24, 30, 36, 45, 60, 75, 90, 105, 120, 150, 180, 210 and 240). The rotation of the platform takes 6 seconds to tilt horizontally. Each experiment included at least 30 repetitions. Experiments were carried out over a range of times during the day, and any circadian contributions (**?**) are expected to be negligible.

### Data collection

As described in Fig. 1a, the local frame of the rotating platform is (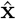, **ŷ**), where **ŷ** is parallel to the upright position of coleoptiles. The angle of the stimulus *θ*_*p*_ is defined as the angle between the vertical of the platform and the direction of gravity, cos *θ*_*p*_(*t*) = **ĝ** ·**ŷ** (*t*). We extracted the center line of each coleoptile, for each image, using a version of the Python-based software Interekt (51). From the center line we extracted the local angle *θ*(*s, t*) of the coleoptile with respect to **ŷ**, the apical and basal angles *θ*_tip_(*t*) = *θ*(*L, t*) and *θ*_base_(*t*) = *θ*(0, *t*), the total length *L*(*t*), and the radius averaged over *s, R*(*t*). The quantities *θ*_tip_ and *θ*_base_ were obtained by locally averaging *θ*(*s*) over 2 mm, i.e. less than *L*_gz_*/*10 where *L*_gz_ is the growth zone. This length scale also sets the minimum required size for a coleoptile to be included in our analysis, i.e. 4 mm. We also calculated *l*⊥(*t*) = ds sin(*θ*(s, t) −*θ*_P_(t)), the projection of the organ perpendicular to ĝ. These quantities were further smoothed with a combination of a median filter and a lowpass filter to remove aberrant data points and allow the computation of smooth time derivatives.

### Numerical simulations

We ran numerical simulations implementing a discretized version of Eq. 2. The code was written in python and based on (12), however allowing to implement any memory kernel or inclination protocol (including transient tilts or clinostat). Required parameters, such as *μ*(*t*), *β, γ, L*, and *θ*(*s, t* = 0), were extracted from experiments (Fig. S5).

### Estimation of gravitropic and proprioceptive gains *β* and *γ*

The estimation of the *β* and *γ* was carried out from the permanent stimulation experiments. In order to compare the dynamics of the growing coleoptiles with our model and simulations, which do not take growth into account explicitly, we consider the normalized gravitropic response defined as Δ(*t*) = *∂*_*t*_ (*θ*_tip_ −*θ*_base_) ·*R/v*_g_, where *R* is the radius of the organ and *v*_g_ = *dL/dt* is the growth rate. The maximal value is termed the dimensionless gravitropic sensitivity 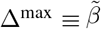 (45, 51, 52). In the limit of negligible growth we have 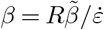 where 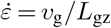 is the average elongation rate of the plant organ and *L*_gz_ the length of its growth zone (42). *γ* is then extracted from the convergence length *L*_c_ = *γ/β*, found by fitting the steady-state angular profile *θ*(*s*) of each organ to an exponential (42): *θ*(*s*) = Θ (1 −exp(−(*s −s*_0_)*/L*_c_)) + *θ*_0_. For each plant, the fit was repeated and averaged over the last 100 min of the experiment. When *s*_0_ *>* 0 we can approximate *L*_gz_ ≈*L −s*_0_. Extracted values are found in Fig. S5 in the SM.

### Memory kernel estimation

In the case of transient inclination, the gravitropic sensitivity *β* is assumed constant, while the inclination angle *θ*_P_(*t*) is time-dependent. Integrating Eq. 2 over *s*, and assuming gravistimulation is zero for *t <* 0, yields: 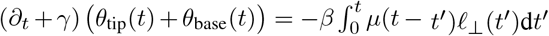, where *l*_⊥_ is the signed length of the organ projected on the axis normal to the stimulation direction. This can then be rewritten as a Volterra integral equation of the first kind on *μ* with *l*_⊥_ as a kernel: 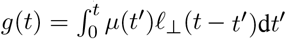, where *g* = *−*1*/β* (*∂*_*t*_ + *γ*) (*θ*_tip_ + *θ*_base_). This integral equation is then solved in the classical way by first turning it into a Volterra integral of the second kind by differentiating it with respect to time, and then discretizing it (53). The initial value is 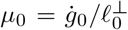, and the solution is then built by successive iterations, for each i: 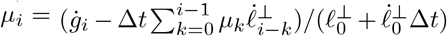. In the case of finite stimuli *τ*_s_, we approximate the sharp transitions of *θ*_*p*_ with a sigmoid curve with characteristic time Δ*t* = 1 min. We have successfully tested this algorithm on a set of Volterra integral equations of the first kind where an analytical solution was known (54, 55). We validated the described method for our model in Eq. 2. We ran simulations for transient inclinations of duration *τ*_s_ with arbitrarily chosen memory kernels *μ*_in_. From the simulated dynamics, memory kernels were estimated using the method described here, and were found to be consistent with the original, regardless of the shape of the kernel or the stimulus duration *τ*_s_ (see Fig. S6). Extracting of memory kernel from experimental data: for each coleoptile we extracted *μ*(*t*) from the trajectory *θ*_tip_(*t*). We discarded instances where the algorithm diverged. We then averaged over repetitions resulting in an average *μ*. We do not take an explicit account of growth when estimating of *μ*, and to the first approximation we neglect the possible slow drift of *l*_⊥_ due to growth.

## AUTHOR CONTRIBUTIONS

Methodology, M.R.; Formal analysis, M.R.; Investigation, M.R. and Y.M.; Writing – Original draft, M.R. and Y.M.; Writing – Review and editing, M.R. and Y.M.; Funding acquisition, Y.M.; Supervision, Y.M.; Conceptualization, Y.M.

## DATA AVAILABILITY

http://dx.doi.org/10.5281/zenodo.8191754

