## Supplementary material for "Plants sum and subtract stimuli over different timescales"

**This PDF file includes:**

**Supplementary Figures**

Fig. S1: Influence of perturbations in $\beta$ and $\gamma$ on memory kernel estimation
Fig. S2: Dependence of memory kernel shape on stimulus duration
Fig. S3: Memory, not relative orientation, is the leading effect in summation and subtraction behaviors
Fig. S4: Dependence of extracted memory kernel on a second kernel in the proprioceptive term
Fig. S5: Estimation of experimental parameters
Fig. S6: Numerical validation of the memory kernel estimation method

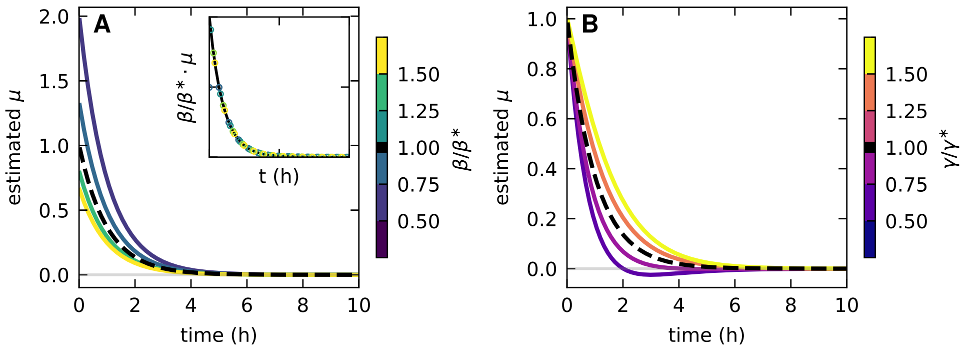


Figure S1: **Influence of perturbations in $\beta$ and $\gamma$ on memory kernel estimation**. We perform simulations (Materials and Methods) of gravitropic responses to transient stimuli, with an arbitrarily chosen exponential memory kernel $\mu^*(t) = A\exp(-t/\tau_{\mathrm{m}})$ and gains $\beta^*$ and $\gamma^*$. From the simulation trajectories $\theta_{\mathrm{tip}}$ we extracted the memory kernel, assuming different values $(\beta, \gamma) \neq (\beta^*, \gamma^*)$. (A) Estimated memory kernels assuming different values of $\beta \neq \beta^*$, compared to the original memory kernel. The estimated memory kernels are multiplied by a coefficient, without affecting the characteristic form. Inset: rescaling the extracted $\mu(t)$ by $\beta/\beta^*$ recovers $\mu^*(t)$. From measurement errors in Fig. S5H, in our case we have $0.67 < \beta/\beta^* < 1.34$, i.e. the perturbations are not significant. (B) Estimated memory kernels assuming different values of $\gamma \neq \gamma^*$, compared to the original memory kernel. From measurement errors in Fig. S5I, in our case we have $0.75 < \gamma/\gamma^* < 1.25$, well within the range of negligible effects.

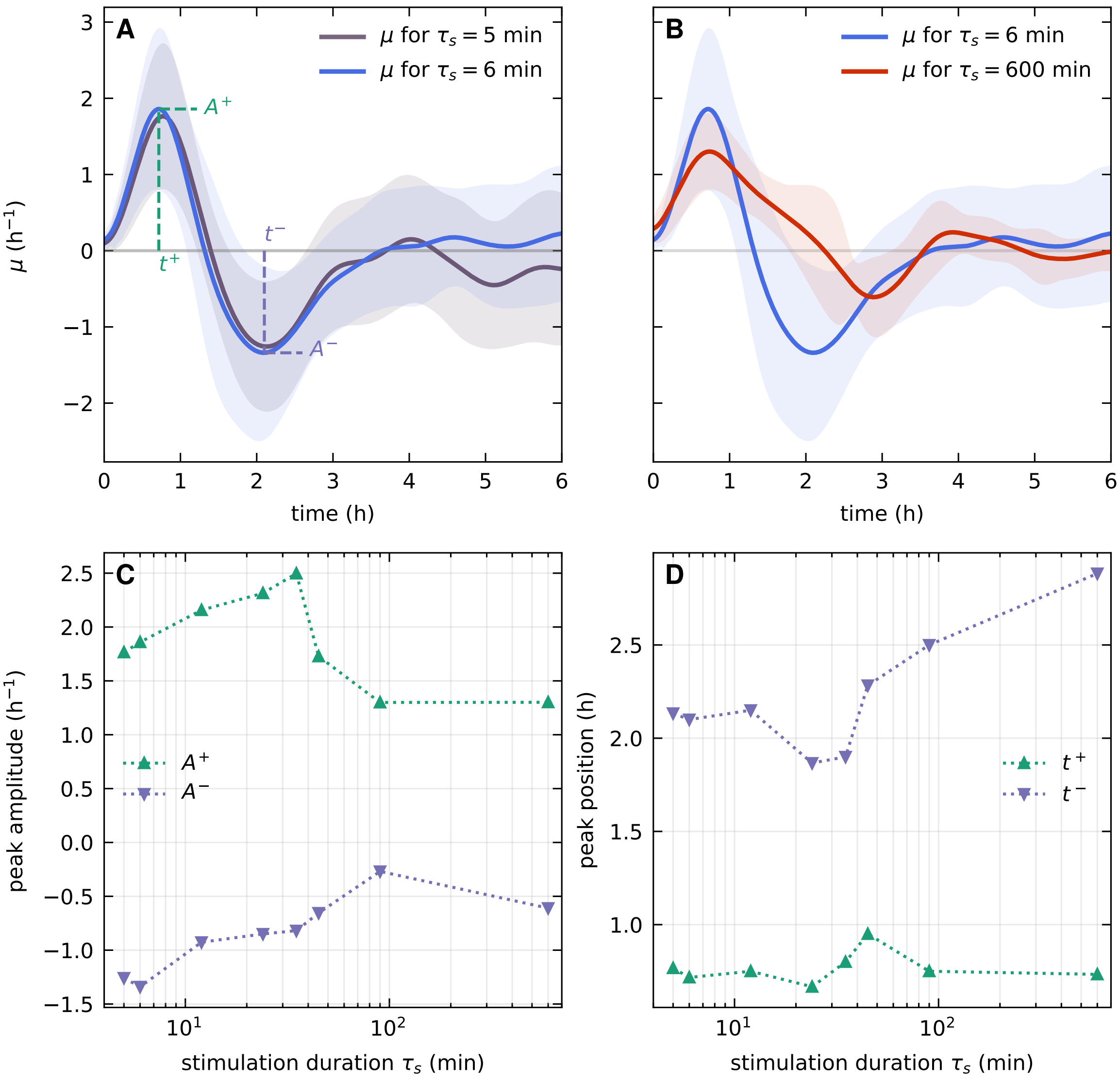


Figure S2: **Dependence of memory kernel shape on stimulus duration.** (A) Comparison of two estimated memory kernels for close values of $\tau_s$ (gray: $\tau_s = 5\,\text{min}$, blue: $\tau_s = 6\,\text{min}$). We define the amplitude of the positive and negative peaks as $A^+$ and $A^-$, and their occurrence times $t^+$ and $t^-$ accordingly. The kernel characteristics are similar, suggesting kernels are robust to perturbations in $\tau_s$. (B) Estimated memory kernels for two limiting values of $\tau_s$: $\tau_s = 6\,\text{min}$ (blue), and $\tau_s = 600\,\text{min}$ (red). The characteristic form of the two kernels are identical (positive peak followed by a negative peak, however the timescale stretches for long stimulation times. (C) Amplitudes of positive peaks $A^+$ (green) and negative peaks $A^-$ (purple) for all extracted memory kernels as a function of stimulation duration $\tau_s$ (Fig. 2C). Amplitudes seem to decrease as $\tau_s$ increase. (D) Peak position, $t^+$ and $t^-$, as a function of $\tau_s$. The position of the positive peak does not seem to depend on $\tau_s$, however the second peak seems to occur later for larger $\tau_s$.

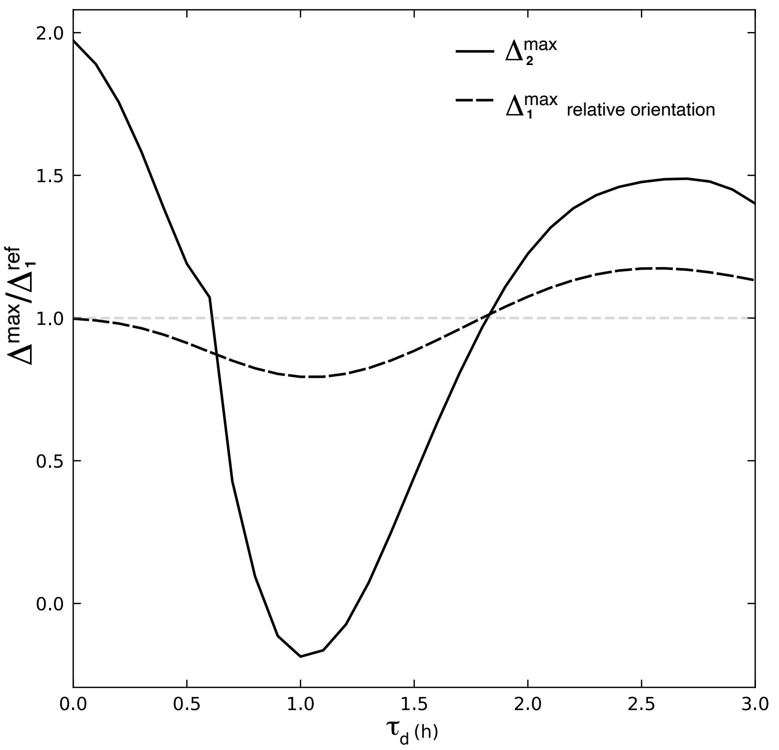


Figure S3: **Memory, not relative orientation, is the leading effect in summation and subtraction behaviours.** We determine whether the summation and subtraction effects found in Fig.3J are a result of the memory kernel we extract, or rather an artifact due to the orientation of the coleoptiles relative to the gravity at the time of the second stimulus. We plot $\Delta_2^{max}$ the maximal simulated response to two stimuli separated by different delay times $\tau_d$, replicating Fig.3J (solid black). In red we plot the maximal simulated response $\Delta_1^{max}$ to a single stimulus $\tau_s = 6$ min, where the initial orientation is taken from the time of the second stimulus in two-stimulus simulations, $\theta_1(s, t = 0) = \theta_2(s = L, t = \tau_s + \tau_d)$. Both responses are normalized by the reference maximal response to a single stimulus with initial straight orientation $\Delta_1^{ref}$. Comparing this response to the response to two stimuli, we find that while the relative orientation does have an effect, the form of the memory kernel is the dominant driver of the observed phenomena.

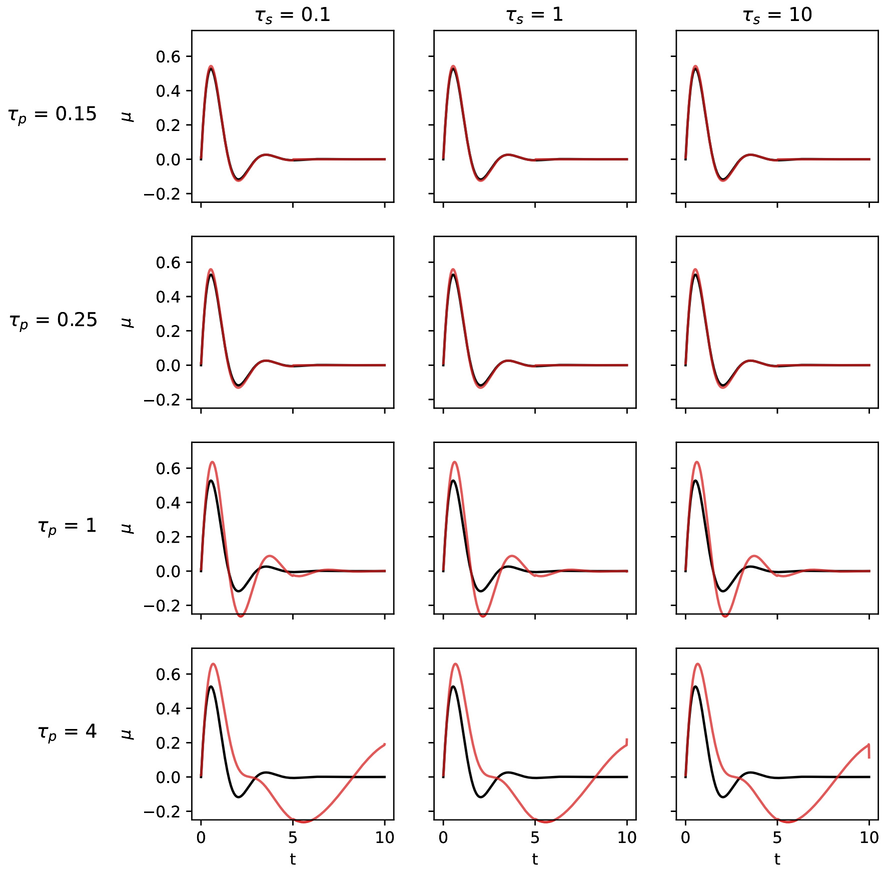


Figure S4: **Dependence of extracted memory kernel on a second kernel in the proprioceptive term.** In our work we assume a memory kernel only in the gravitropic term. While the processes underlying proprioception are unknown, it is possible that the proprioceptive term $\gamma\kappa(s,t)$ may also be described with a memory kernel, $\gamma\int_{-\infty}^{t}\mu_P(t-t')\kappa(s,t')\mathrm{d}t'$. Here we assess the effects of such a (neglected) kernel on the form of the estimated gravitropic memory kernel. We ran simulations with different stimulus times $\tau_s = \{0.1, 1, 10\}$, and a different kernels in the proprioceptive term. The input kernels followed the form $\mu(t) = \exp(-t/\tau)\sin(2\pi/3t/\tau)$, with $\tau = 1$ for gravity and $\tau = \{0.15, 0.25, 1, 4\}$ for proprioception, and normalized. From the trajectories we extracted the the memory kernel associated with gravitropism, as done in the main text, neglecting the underlying proprioception kernel. The extracted gravitropic kernel (red) is plotted with the input gravitropic kernel (black) for comparison. We find that effects become considerable for proprioceptive kernels with timescales $\tau$ significantly larger than the gravitropic kernel, which is physiologically unlikely since growth is the limiting timescale. We also find that a kernel in the proprioceptive term does not explain the observed dependence of $\mu$ on $\tau_s$.

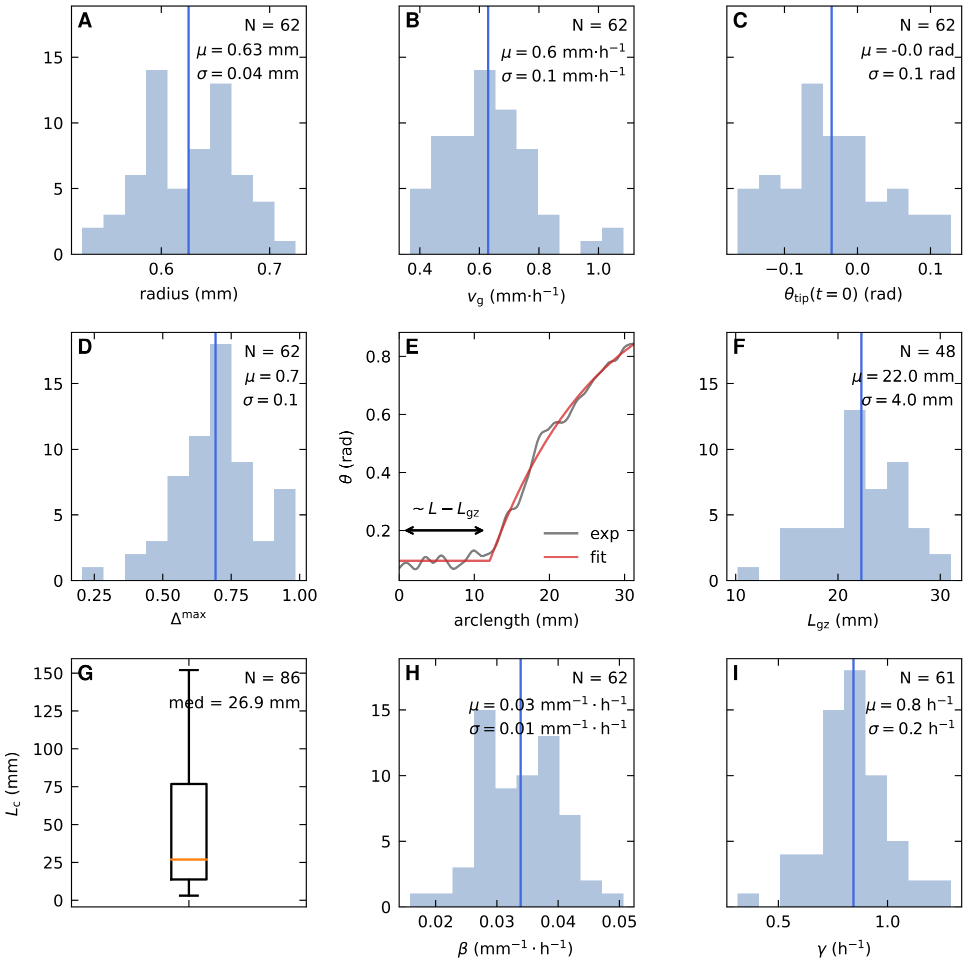


Figure S5: **Estimation of experimental parameters.** (A) Distribution of coleoptile radii averaged over the experiment duration. (B) Distribution of the average growth velocity $v_g$. (C) Distribution of the initial tip angle $\theta_{\text{tip}}(t = 0)$. (D) Distribution of the maximum gravitropic response $\Delta^{\text{max}}$. (E) Steady-state orientation profile of a coleoptile (gray) and fit to the predicted exponential profile (*1*) (red). The constant part roughly corresponds to the non-growing region the coleoptile. This fit outputs $L_{\text{gz}}$ and $L_c$. (F) Distribution of the estimated growth zone length $L_{\text{gz}}$. Consistent with (*2*). (G) Distribution of convergence length $L_c$. Outliers not shown. This representation was chosen because of the heavy tail of the distribution.(H) Distribution of the estimated gravitropic sensitivity $\beta$. (I) Distribution of the proprioceptive sensitivity $\gamma$. Vertical blue lines: $\mu$=avg; $\sigma$=std.

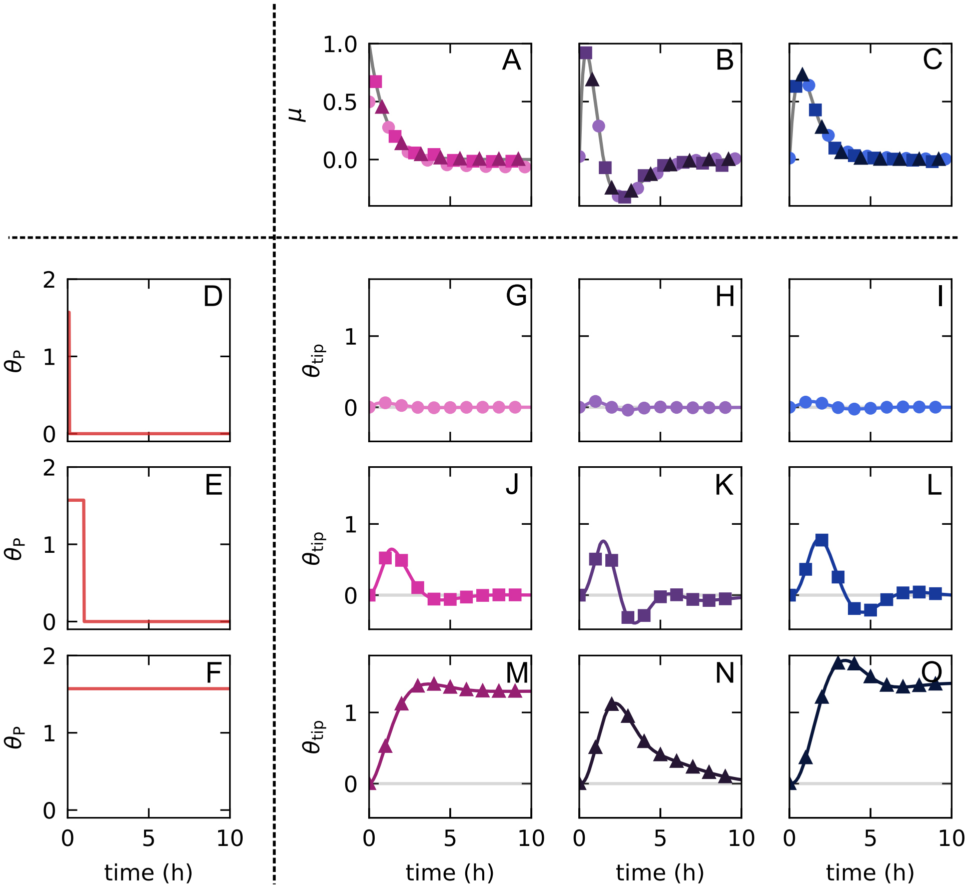


Figure S6: **Numerical validation of the memory kernel estimation method**. (A–C) Simulations were run with different input memory kernels (columns, from left to right): $\mu(t) = A\exp(-t/\tau_A)$ (pink), $\mu(t) = B\exp(-t/\tau_B)\,(b_0(t/\tau_B) - b_2(t/\tau_B)^2)$ (purple), $\mu(t) = C\exp(-t/\tau_C)\,(c_0(t/\tau_C) + c_2(t/\tau_C)^2)$ (blue); (D–F) and with different stimulation durations (rows, from top to bottom): $\tau_s = 10dt$ (light circles), $\tau_s = 1/\gamma$ (neutral squares) and $\tau_s = 10\,\text{h}$ (dark triangles). (G–O) For each combination of $\tau_s$ and input $\mu$, we obtain the dynamics of the tip angle $\theta_{tip}(t)$. (P–R) Estimated memory kernels, extracted from each $\theta_{tip}$ previously obtained. Note how, for each input $\mu$, the kernels estimated from different $\tau_s$ (different symbols) collapse on the curve of $\mu_{input}$ (grey).
